# LSTM-Attention-Guided Graph Neural Networks for Integrated Genotype–Environment Modeling in Maize Yield Prediction

**DOI:** 10.1101/2025.11.09.687529

**Authors:** Amir Morshedian, Mike Domaratzki

## Abstract

We present a deep-learning framework that combines an LSTM, a graph neural network (GNN), and transformer-style attention to model genotype–environment (G × E) effects for maize yield prediction. Weather data for a growing season is summarized using LSTM and encoded into a 21-dimensional embedding that is used as the environment node feature; 437,214 SNPs are summarized into 548 principal components that instantiate genotype nodes. Multi-head attention dynamically weights the edges during message passing. We compare three architectures: A (fully bipartite graph), B (A with intra-set top-*k* similarity within genotype and within environment), and C (B with a single learnable supernode readout that attends over all nodes after message passing). The joint representations feed a compact MLP for yield prediction. Using a forward-time split (2014–2021 train; 2022 test with unseen genotypes and unseen environments), performance improves monotonically from A to C: A (RMSE 2.7749, PCC 0.4115, R^2^ 0.1693), B (2.3683, 0.6622, 0.4385), C (2.2120, 0.6945, 0.4823). Compared to A, C has a reduction in RMSE by 0.5629 (∼20.3%) and an increase in PCC by 0.283 (∼68.8%), indicating that global, content-adaptive aggregation promotes local G *×* E propagation. Performance of our approach remains consistent regardless of the number of genotypes per environment and has strong performance under variable or unbalanced genotype sampling expression across environments.

## 1 Introduction

Global food security critically depends on functioning agricultural systems. However, it is being compromised under the pressures of increasing global population and accelerating climate change, characterized by more frequent and severe droughts and heat waves. The United Nations projects the world population to reach 9.7 billion by 2050 [1]. Additionally, 2023 was the warmest year on record [2]. Harvest data have long been the standard benchmark for evaluating crop performance. Yet, a key limitation is that year-to-year variability in environmental drivers makes such records insufficient for predicting the performance of future crops under changing conditions [3, 4]. Therefore, farmers and policymakers require effective and rapid methods to assess yield at the regional scale to enable pre-allocation of inputs and informed risk management decisions.

With the advancement of next-generation sequencing technologies, dense genome-wide markers (SNPs) can now be collected cost-effectively, allowing genomic selection to estimate breeding values from genomic data only and accelerate genetic gain [5]. Through estimates of allele sharing between individuals, genomic markers allow understanding of the relationship between varieties, thus providing informative genotypic features for predicting agronomic performance [6].

The environment in which a particular variety is grown is also important. The field crop environment comprises atmospheric conditions, soil properties and management practices, including precipitation, temperature, solar radiation, wind, humidity, photoperiod, soil electrical conductivity and pH, as well as practices such as planting patterns, irrigation, fertilizer applications and pest management. Because these drivers vary between seasons and locations, models that do not account for environmental variability are likely to not accurately predict yield in all situations and perform poorly under extreme climatic conditions [7].

Yield emerges from genotype-by-environment (G × E) interactions, a phenomenon whereby varieties producing desired quantitative trait values in one environment may fail to provide the same outcomes in another environment [8]. It is therefore beneficial to model both genomic markers and environmental variables simultaneously when predicting yield. Consequently, any tools developed to aid crop breeding may benefit from replicating the impact of G×E by incorporating both genetic and environmental information.

Many solutions have been proposed to address crop yield prediction using G × E interactions [10, 16, 18, 23, 25, 29]; however, a substantial research gap still exists in effectively capturing their interaction. Most existing methods struggle to encode environmental features for growing season and we expand their approach to apply to unseen genotypes and unseen environments. Another major challenge lies in modeling large-scale datasets with thousands of genotype and environment records, robust feature representation.

In this paper, we propose a G × E modeling framework that encodes daily weather data using long short-term memory networks (LSTM) and genotypes using principal component analysis (PCA) into compact embeddings. Subsequently using both through a graph neural network with multi-head attention to learn G × E interactions. We presented three graph architectures: A (fully bipartite graph), B (A with intra-set top-k similarity within genotype and within environment), and C (B with a single learnable supernode for readout). The framework is assessed under a forward-time train/test split with unseen genotypes and environments. Results demonstrated a consistent improvement from A to C and robustness to uneven genotype sampling per environment. We compare our results between all three architectures with the G2F G × E Competition results [9]. The primary contributions of this manuscript are as follows:

1. Unique environmental representation: We present a compact representation of the season-long harvest and the related weather data using LSTM allows the model to use environmental context better than the previous approaches.
2. Progressive attention-weighted graph densification: We start with a GNN baseline to model G × E interactions and then we increase the edge density systematically to determine out the impact of the graph connectivity and the model complexity on the accuracy of the yield prediction.

## 2 Literature review

Several approaches have been proposed for crop yield prediction, focusing on G × E interactions, ranging from statistical models to advanced deep learning methods.

Genomic best linear unbiased prediction (GBLUP) is a linear mixed model for yield prediction in multi-environment trials [10, 11, 12], and can be extended to capture genotype × environment (G × E) interactions. As an example, Jarquín et al. [10] introduced a reaction norm model where genetic and environmental gradients are described as linear functions of markers and environmental covariates, respectively. Their method used covariance structures to model interactions between high-dimensional sets of markers and environmental covariates.

A limitation of linear models such as GBLUP is that they capture only linear relationships between genotype and environment, often failing to represent complex non-linear genotype × environment interactions [13]. To better capture high-order G × E interactions, researchers have therefore shifted to the use of nonlinear machine-learning methods.

Tree-based ensemble methods such as random forests and gradient boosting have been used to yield prediction problem [14, 15, 16]. For example Fernandes et al. [16] used LightGBM and tree-based feature partitioning to model non-linear G×E interactions.

Deep learning methodologies have been introduced that can handle the high dimensionality of input data and capture complex, nonlinear feature interactions. These models are increasingly applied in genomic selection and crop yield prediction within agricultural research. Convolutional neural networks (CNNs) are one from of deep learning used to model G × E for crop-yield prediction [17, 18, 19].

Feng et al. [18] presented a CNN-based based approach that processes environmental data through a CNN branch to capture temporal and spatial patterns, while genotypic marker data is fed into a separate CNN to extract genomic features. These two feature representations are then merged in a joint layer. The combined features are passed through another MLP as a predictor for yield prediction.

Recurrent neural networks, in particular, long short-term memory (LSTM) networks, have been preferred for modeling time series weather data and other sequential inputs [20]. LSTMs suitably model long-term dependencies in climate sequences in a growing season that are very important for understanding how drought periods or temperature fluctuations at different growth stages affect final yield. Zhong et al. [21] employed a spatio-temporal deep learning framework based on LSTM, integrating remote sensing and climate data to predict maize yield and detect extreme yield losses. However, this approach used only environmental features.

Attention mechanisms have been also implemented on RNNs to help determine the most effective time windows or features for forecasting [22]. Shook et al. [23] proposed an LSTM–RNN with attention that combined genotype clusters and weather time-series to predict maize yield. The LSTM captured temporal patterns, while the attention layer highlighted the most important periods of the growing season, improving the model’s ability to represent G×E interactions. CNN-LSTM hybrid models have also been utilized for spatial and temporal pattern recognition [24].

Yao et al. [25] proposed GEFormer, a G × E interaction-based genomic prediction method that used both genomic and environmental data in a unified architecture. It employed a MLP to capture global and local patterns in genomic marker data. To handle temporal dependencies in weather and other environmental factors, GEFormer incorporated a linear attention mechanism, which improved the model’s ability to focus on critical time windows.

In this paper, we adopt GNNs for modelling G × E. GNNs have previously been adopted for genomic selection without environmental data. He et al. [26] introduced HGATGS, a graph attention network designed for genomic selection. The method represented individuals as nodes connected by edges built from genomic similarity, enabling the model to exploit higher-order relationships among genotypes. Through graph convolution and attention mechanisms, HGATGS adaptively weights genetic contributions. Kihlman et al. [27] proposed sub-sampling graph neural networks (GCN-RS) for genomic prediction of quantitative traits. They constructed graphs that represent genomic relationships among individuals and applied graph convolutional layers to propagate marker information. To improve scalability, the sub-sampling strategy reduces graph size while retaining key genetic structure, allowing efficient training on large datasets. Both of these graph-based approaches rely only on genotype sequences and do not incorporate environmental features.

Transformers are another deep learning method used in genomic prediction and yield prediction [28, 29]. As an example, Zou et al. [29] assembled a large-scale multi-environment dataset wheat yield records, paired with real-valued environmental data. They propose a multimodal deep learning framework that uses both Bi-LSTM and Transformer architectures to model temporal environmental variables. Genotype data are encoded in a parallel branch, and the learned genotype and environment embeddings are fused for trait prediction.

## 3 Material and Methods

### 3.1 Graph Foundations

Graphs are used to model complex and interconnected systems in the real world, such as knowledge graphs, social networks, and molecular structures. Formally, a graph *G* = (*A, H*) is a set *V* of *n* nodes connected by edges. *A* denotes an *n × n* adjacency matrix where each entry *a*_*ij*_ ∈ {0, 1} indicates the presence or absence of an edge connecting nodes *i* and *j*. Additionally, we define *N*_*i*_ as the set of neighbors of node *i*, which are the nodes connected to *i* by an edge, i.e., *N*_*i*_ = { *j* ∈ *V* | *a*_*ij*_ = 1 }.

### 3.2 Message-Passing Graph Neural Networks

Graph Neural Networks (GNNs) are a class of deep learning architectures designed to operate on graph-structured data. GNNs leverage the graph topology to propagate and aggregate information between connected nodes. In a GNN, each node is associated with a feature or attribute vector that encodes its initial information. Accordingly, the matrix of initial embeddings **H** ∈ ℝ^*n×d*^ stores these attributes, where each row **h**_*i*_ ∈ ℝ^*d*^ represents the features of node *i* [30]. For example, in molecular graphs, each node contains information about the atom type, and edges can represent interactions among atoms. They follow the principle of message passing, where each node iteratively updates its embeddings by aggregating information from its local neighbors [31]. Formally, the node embeddings 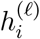 for each node *i* ∈ *V* is updated from layer *ℓ* to *ℓ* + 1 via a two-step message-passing process:

**(1) Message construction:** For each node *i* and its neighbors *j* ∈ *N*_*i*_, construct a message 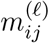 that captures the relationship between the embeddings of nodes *i* and *j*.

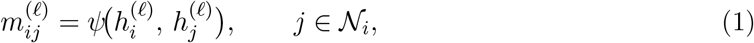

with a function *ψ* that can be any problem-appropriate function to construct edge messages.

**(2) Neighbour aggregation:** Combine all messages from the neighbors of node *i* to produce a single aggregated message 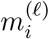:

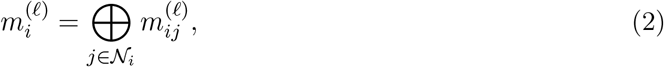

where ⊕ is an operator (e.g., sum, mean, or max) that aggregates messages from all neighbors *j* ∈ *N*_*i*_, and finally the node embedding is updated:

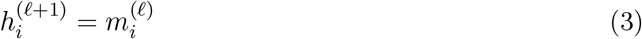

### 3.3 Prediction from embeddings

Initially every node is assigned its raw embedding. Equations (1)–(3) iteratively refine these embeddings, 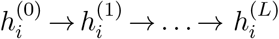, incorporating increasingly larger neighbourhood context. The final embedding matrix 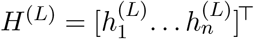 serves as input predictor:

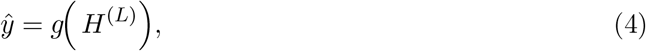

where *g* denotes a predictor such as a linear classifier, bayesian regression head or multilayer perceptron.

### 3.4 Graph Attention Network

A class of GNNs uses attention mechanisms to weight the importance of different neighbors during aggregation [32]. Graph attention network refine (1)–(2) by assigning *learned, data dependent* weights to each neighbour:

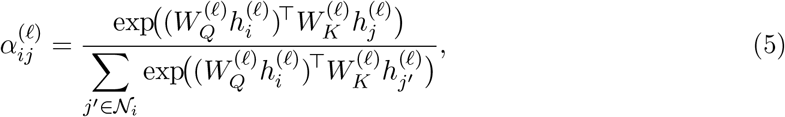

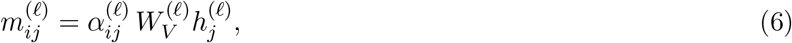

where 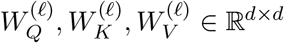 are learned projections. The node update thus becomes

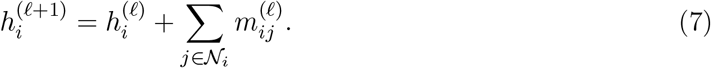

We will employ Graph Attention Networks (GATs) as our main model to represent G × E in Section 3.8.

### 3.5 Dataset

We have utilized the Genomes to Fields (G2F) Genotype × Environment Prediction Competition data [9], which is a large multi-year maize field trial resource and is specifically designed to benchmark G×E models. The 2023 release has over 180,000 individual field plots which are assessed under nearly 280 unique year-location combinations in North America. In total, the experiment comprised 4,683 distinct hybrids in the training dataset, while 4,928 hybrids are present in the full genotypic dataset.

#### 3.5.1 Phenotypic Data

Every field plot has central agronomic characteristics such as grain yield (converted to 15.5% moisture, given in Mg ha^−1^), days to anthesis, days to silking, and the anthesis-silking interval (ASI). The trials were performed according to standard management practices, and the majority of hybrids were replicated in several plots in the environments. The summary statistics indicate that considerable variability exists not only between environments (like the differences in the average yield from one site to another) but also within environments (the variation among replicates), making it a realistic challenge for modelling.

#### 3.5.2 Genotypic Data

Through the use of publicly available parental marker profiles, hybrid genotypes were deduced. The final marker matrix consists of 437,214 high-quality SNPs (Single Nucleotide Polymorphisms) coded as 0, 1, or −1 that were recorded in the 4,928 distinct hybrids. The filtering process eliminated SNPs with a minor allele frequency below 0.01 or a missingness above 5%.

#### 3.5.3 Weather Data

The weather data were obtained from the G2F competition dataset [9], which provides daily meteorological records for each site-year pair. These records were originally retrieved from the NASA POWER archive and include radiation variables (All Sky Surface PAR Total, All Sky Surface Shortwave Downward Irradiance, and All Sky Surface Shortwave Downward Direct Normal Irradiance), temperature measures (wet bulb, maximum, minimum, and mean at 2 m), humidity indicators (specific and relative at 2 m), dew/frost point (2 m), precipitation corrected, surface pressure, wind speed (2 m), and soil moisture metrics at multiple depths (profile soil moisture, surface soil wetness, and root zone soil wetness). We standardized all features to zero mean and unit variance within each environment before modeling to ensure comparability across sites and seasons.

#### 3.5.4 Cross-Validation and Data Splitting

In order to imitate actual breeding, a forward-time validation strategy was employed which involved the training of all models on the data from the field trials carried out from the year 2014 to 2021. The 2022 trials, on the other hand, were kept as a test set that was not used for training the entire model as unseen genotypes and unseen environments. During the 2014–2021 period, we monitored that no model observed the same combination (Genotype-Environment) in both training and validation, thus providing an estimate of performance on truly unseen contexts. We used 80% for training and 20% for validation. The 2022 trials, which were not used at any point during the modeling period, are used only on the final evaluation which shows the ability of each model to apply the learned G×E relationships to a new season with new genotypes.

### 3.6 Genotype Dimensionality Reduction

In order to cope with the high dimensionality of our genotype dataset (4,928 samples by 437,214 SNPs), we applied a two-step reduction strategy. First, we performed probabilistic imputation to replace missing entries: for each SNP column, we estimated the empirical frequencies of {− 1, 0, +1 } and stochastically sampled replacements according to these probabilities. This approach preserves each marker’s marginal distribution without imposing strong assumptions on linkage structure. Subsequently, we used classic variance-based and redundancy filters to remove uninformative or SNPs with very high correlation:

1. Low-Variance Filter: We eliminated all SNP columns that had a single genotype value in at least 95% of the observations.
2. Windowed Similarity Pruning: We moved a window of width *L* along the rest of the markers, calculating the similarity among pairs inside every window. In cases when two SNPs were found to be identical in the samples by at least 95%, we kept just one representative marker; thus, local redundancy was removed while regional haplotype structure was kept intact.

These filters reduced the feature set from 437,214 to 133,673 SNPs. The second stage consisted of performing principal component analysis (PCA) on this filtered matrix. PCA is the identification of orthogonal axes with the most variance which automatically compresses correlated markers to a small number of components. The 548 principal components that we kept, resulted in over 90% of the total genetic variance being accounted for. The resulting 548-dimensional vector provides a compact yet informative summary of each genotype, lowering computational demands and mitigating overfitting risks associated with excessively long input sequences.

### 3.7 Weather-to-Harvest Feature Extraction using LSTM

Among the weather variables mentioned in section 3.5.3 we have kept five variables: All–Sky Surface PAR Total, 2 m Maximum Temperature, 2 m Minimum Temperature, Corrected Precipitation, and All–Sky Surface Short-Wave Direct-Normal Irradiance. This small subset was chosen to span the range of environmental conditions. Accordingly, for each environment we filtered a k*×*5 values, where k denotes the number of growing-season days. As the environmental dataset is large, an effective summarization method is required. Previous approaches have relied on averaging (weekly or monthly) techniques [33]; however, these often conceal short but high-impact events, such as sudden heatwaves during tasseling. To address this, we instead encode the sequence of daily observations using a single-layer unidirectional LSTM.

After *z*-score normalising every feature, the daily vector **x**_*t*_ ∈ ℝ^5^ is fed to the LSTM, whose gating equations are

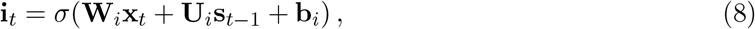

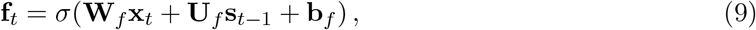

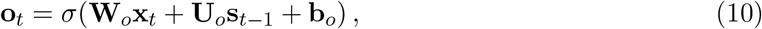

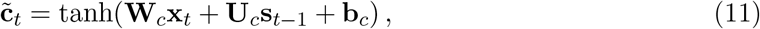

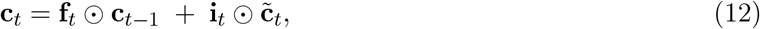

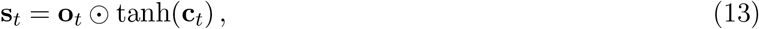

where *σ*(·) is the logistic sigmoid and ⊙ denotes element-wise multiplication [34, 35]. Here, *W*_*i*_, *W*_*f*_, *W*_*o*_, *W*_*c*_ ∈ ℝ^*m×*5^ are the input weight matrices, *U*_*i*_, *U*_*f*_, *U*_*o*_, *U*_*c*_ ∈ ℝ^*m×m*^ are the recurrent weight matrices, and *b*_*i*_, *b*_*f*_, *b*_*o*_, *b*_*c*_ ∈ ℝ^*m*^ are bias terms, where *m* denotes environmental embedding vector length. All weight matrices and biases are learnable parameters optimized during training. At the beginning of each sequence, both the hidden state **s**_0_ and the cell state **c**_0_ are initialized to zero vectors of dimension *m*, and are then updated recursively according to Equations (8)–(13). The final hidden state **s**_*k*_ ∈ ℝ^*m*^, which encodes the entire sequence of *k* daily records, is used as the environmental embedding **z**_env_.

To choose the embedding length we considered values of *m* ∈ {6, 9, 12, 15, 18, 21, 24, 27, 30, 35, 40, 45, 50 }. For each embedding size *m*, the environmental representation **z**_env_ was used to predict yields across all genotype–environment pairs in the validation set, and performance was evaluated using the Pearson correlation coefficient (PCC) between predicted and observed yields. Figure 1 shows that performance peaks at *m* = 21 (PCC ≈ 0.75) using architecture A (described below in Section 3.8.1), plateaus for *m* = 24–27, and then generally declines once *m* ≥ 30. We therefore fix *m* = 21 for all downstream genotype environment fusion models [22], giving us a compact but information-rich representation that captures normal behavior of weather data during the growing season [36].

**Figure 1:**
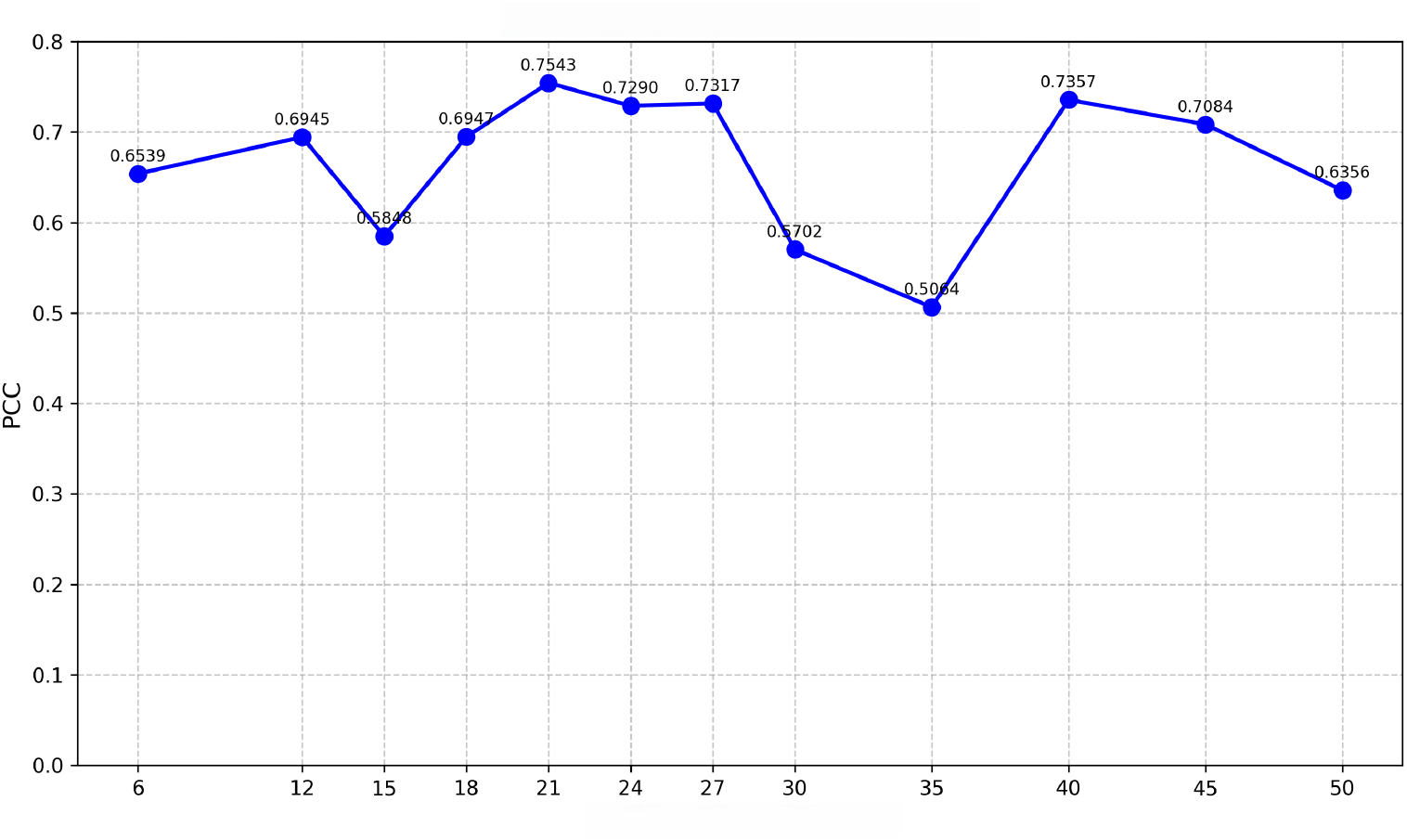
Pearson correlation coefficient (PCC) versus environment vector size *m*. The optimum occurs at *m* = 21.

### 3.8 End-to-End G×E Pipeline

Figure 2 presents the full genotype-by-environment (G × E) pipeline used for crop yield prediction. This figure brings together the individual components introduced in the sections 3.6 and 3.7 into a unified framework. Two complementary embeddings are first derived:

**Figure 2:**
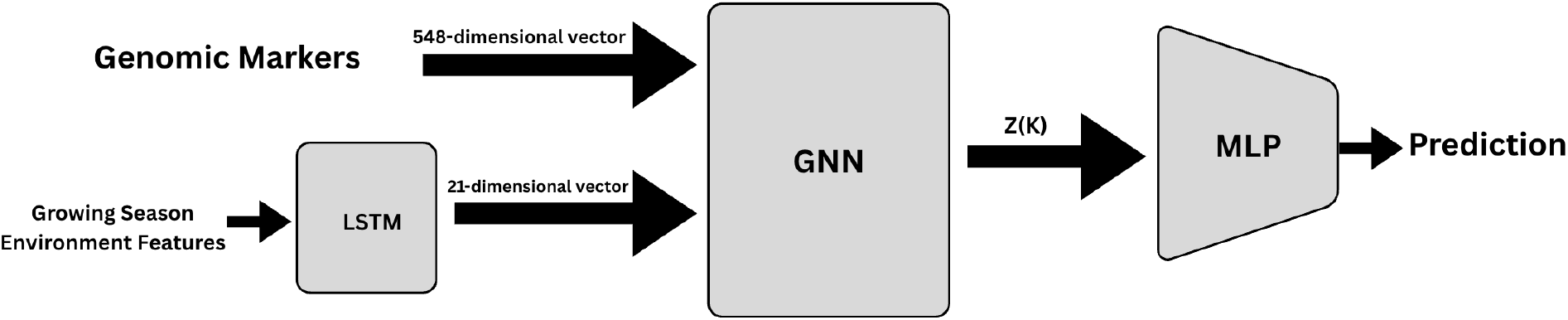
Overview of the genotype-by-environment (G*×*E) pipeline for crop-yield prediction.

- **Genomic stream**: After completing the quality control and imputation processes, 437,214 SNPs are embedded to 548 principal components. The resulting output is a dense marker vector which encodes the major axes of the genetic variation.
- **Weather data set**: The single-layer LSTM reads five variables on a daily basis, i.e. total PAR, maximum temperature, minimum temperature, corrected precipitation, and short-wave DNI. The last hidden state **s**_*k*_ ∈ ℝ^21^, representing the 21-dimensional order-aware weather fingerprint, is denoted as **z**_env_.

Both embeddings are given as node features to a GNN with 21 + 548 = 569 nodes. After *K* propagation steps, we obtain updated node embeddings. The set of embeddings **Z**(*K*) are passed to the multilayer perceptron (MLP) model. **Z**(*K*) in Architectures A and B, consists of the genotype related nodes embeddings, while in Architecture C it corresponds to the supernode embedding. Further details of these architectures are provided in Sections 3.8.1–3.8.3.

The MLP maps the input embeddings to a single scalar value representing the predicted yield. The training process is based on the minimization of mean-squared error with early stopping, while hyper-parameters are selected using the forward-time cross-validation procedure (2014-2021 training seasons, 2022 reserved).

Next, we present three variations of GNN architecture, all of which have the same interconnecting pipeline components as the basic model.

#### 3.8.1 Architecture A

Architecture A represents the interactions between genotype and environment using a fully connected bipartite graph (Figure 3), which is comprised of two distinct node sets.

**Figure 3:**
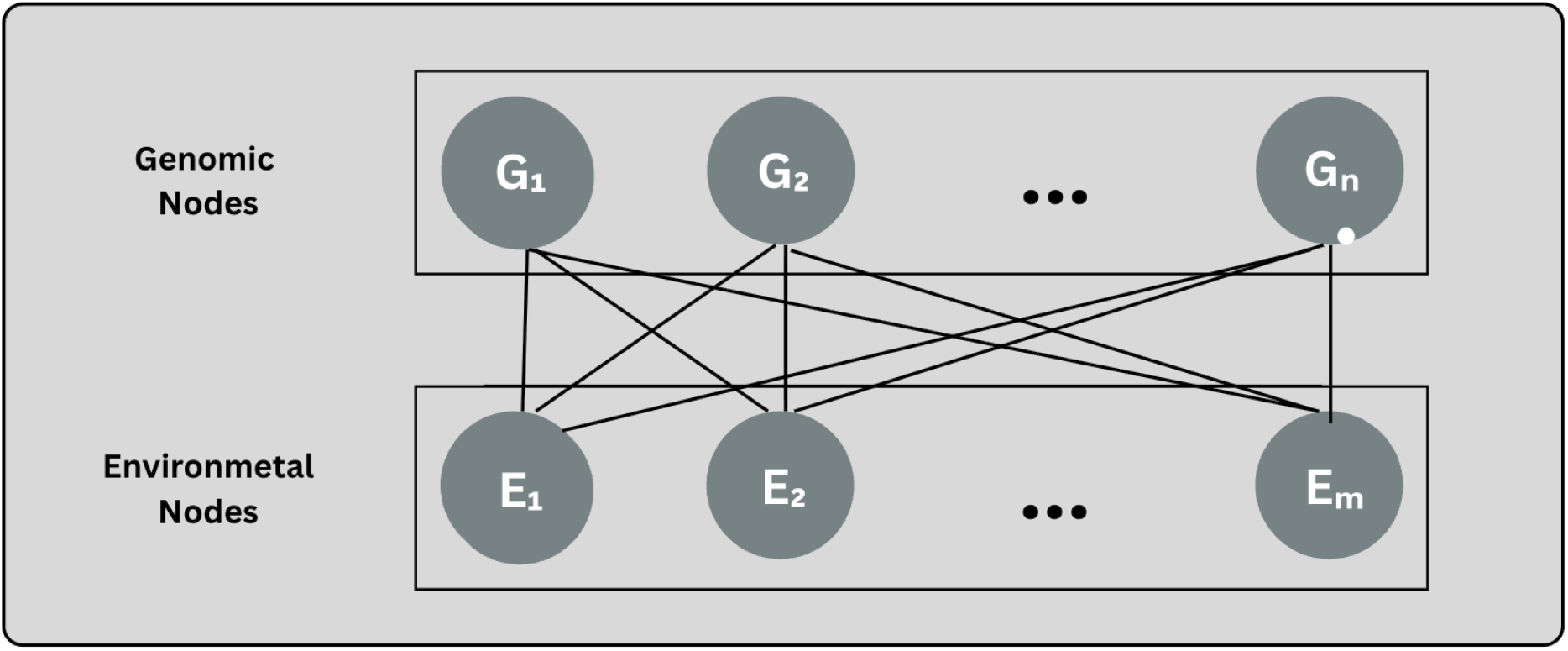
Detailed illustration of Architecture A as a fully connected bipartite graph for modelling genotype–environment interactions with attention.

- **Genomic nodes**: The *n* = 548 principal components scores from the SNP matrix are instantiated as individual nodes *G*_*i*_.
- **Environmental nodes**: The LSTM’s *m* = 21-dimensional weather fingerprint is decomposed into *m* single-feature nodes *E*_*j*_.

Each genomic node connects with each environmental node, which results in *n* × *m* edges. Communication by means of messages across each edge is done through the use of multi-head transformer attention:

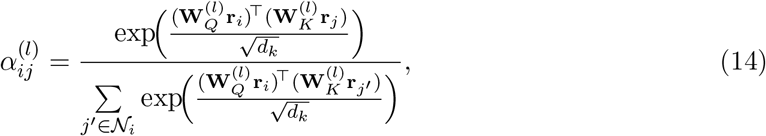

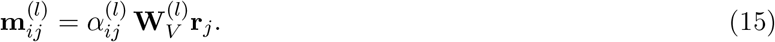

where **r**_*i*_ and **r**_*j*_ are the current embeddings of nodes *G*_*i*_ and *E*_*j*_, and 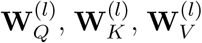 are head-specific projection matrices. Messages from all heads are concatenated and aggregated at the target node; residual connections and layer normalization follow each attention block to stabilize training.

When *K* such propagation layers are stacked together it leads to genomic embeddings which incorporate the total range of genotype–environment interactions evident through the bipartite attention mechanism. Architecture A represents our baseline maximal-capacity design; the following variations will either prune or re-weight the edges and accordingly, explore accuracy–efficiency trade-offs without changing any other pipeline components.

#### 3.8.2 Architecture B

The extension of the bipartite attention framework with Architecture B is done by including intra-set connectivity on both the genotype and environment sides while still keeping all *n* × *m* genotype–environment edges (Figure 4). This improves the information exchange between the two modalities and within each modality, allowing locally conducive embeddings to exchange information during cross-set propagation.

**Figure 4:**
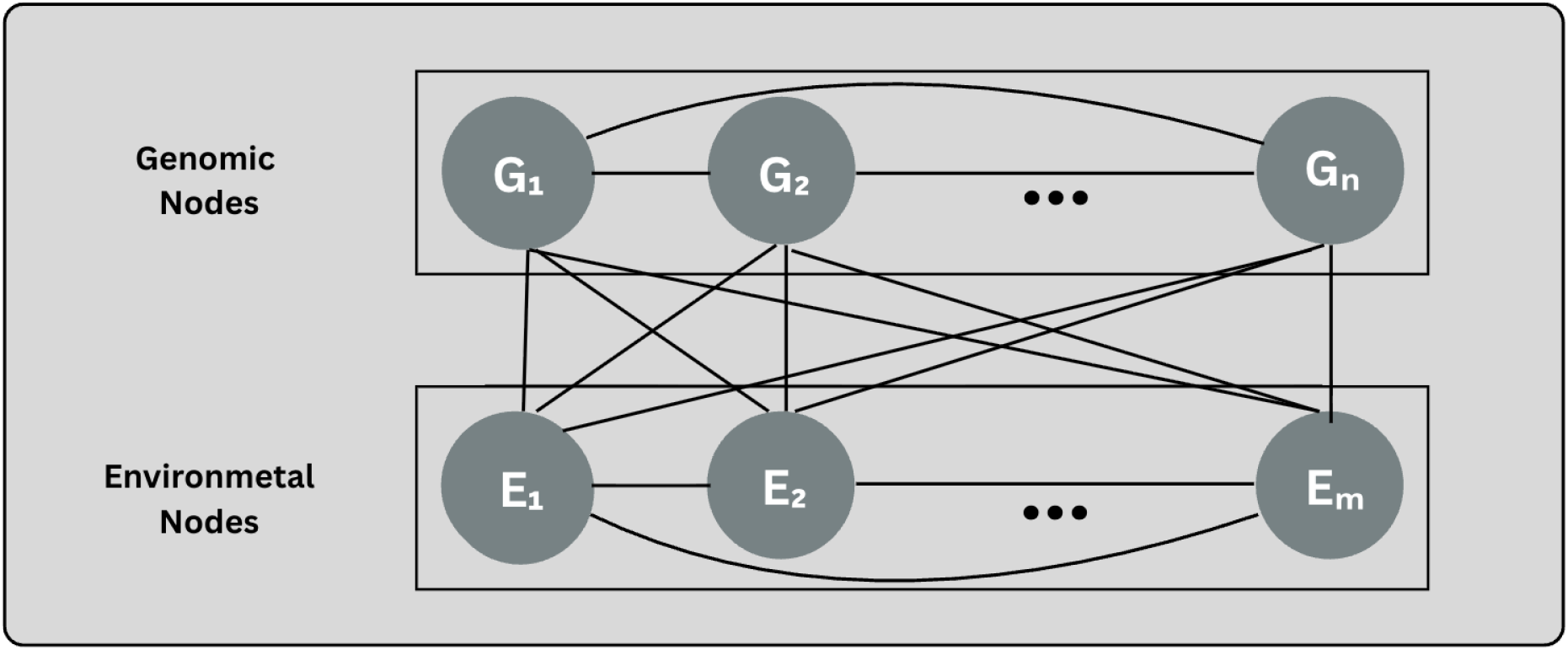
Detailed illustration of Architecture B showing the bipartite genotype–environment structure augmented with intra-genotype and intra-environment similarity edges.

The *n* = 548 PCA components that are extracted from the SNP matrix represent the genomic nodes *G*_*i*_. Each *G*_*i*_ remains connected to the environmental nodes, and in addition to this, each node *G*_*i*_ is attached to its top-*k* most nearest genomic neighbors (*k* = 10) according to

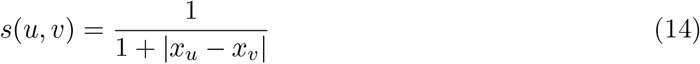

where *u* and *v* refer to individual nodes, and *x*_*u*_ and *x*_*v*_ denote their associated embedding vectors.

The *m* = 21 single-feature weather nodes *E*_*j*_ also conserve their full bipartite links to genomic nodes while each *E*_*j*_ forms the top-*k* most similar environment neighbors (*k* = 10) using equation 14, which thus generates a intra-environment adjacency with per-node out-degree *k*.

Let *A*_*GG*_ and *A*_*EE*_ symbolize the directed intra-set adjacencies (bounded out-degree *k* per node), and *A*_*GE*_, *A*_*EG*_ the bipartite adjacencies between *G* and *E*. The complete message-passing graph is symbolized as

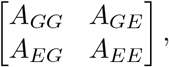

where all of the *n* × *m* bipartite edges of architecture A is present, and the intra-set edges are additionally included.

Message passing is accomplished with the aid of multi-head attention operating over all incident edges. Attention coefficients are computed on any edge (be it intra-set or cross-set), and messages are aggregated through residual connections and layer normalization after each block (as in architecture A). The extra connectivity allows for mixed multi-hop paths; for example, *G*_1_ → *G*_2_ (intra-genotype) and *G*_2_ → *E*_1_ (bipartite) mean that at the end of two propagations, *E*_1_ has obtained information from *G*_1_. More generally, after *ℓ* propagations, a node’s embedding brings together all members of its *ℓ*-hop neighborhood according to the block adjacency, using both intra-set smoothing and inter-set interaction.

This approach captures a similarity-driven local structure while also keeping the full genotype– environment map from the architecture A.

#### 3.8.3 Architecture C

Architecture C changes the graph-level readout, which is the same as the single block-graph, node-level G × E message passing of Architecture B (intra-genotype and intra-environment edges plus full bipartite *G* ↔ *E*). We replace the forwarding of all genomic node embeddings to the MLP with a single learnable supernode query that attends to all nodes and aggregates them into one global embedding via multi-head attention (Figure 5).

**Figure 5:**
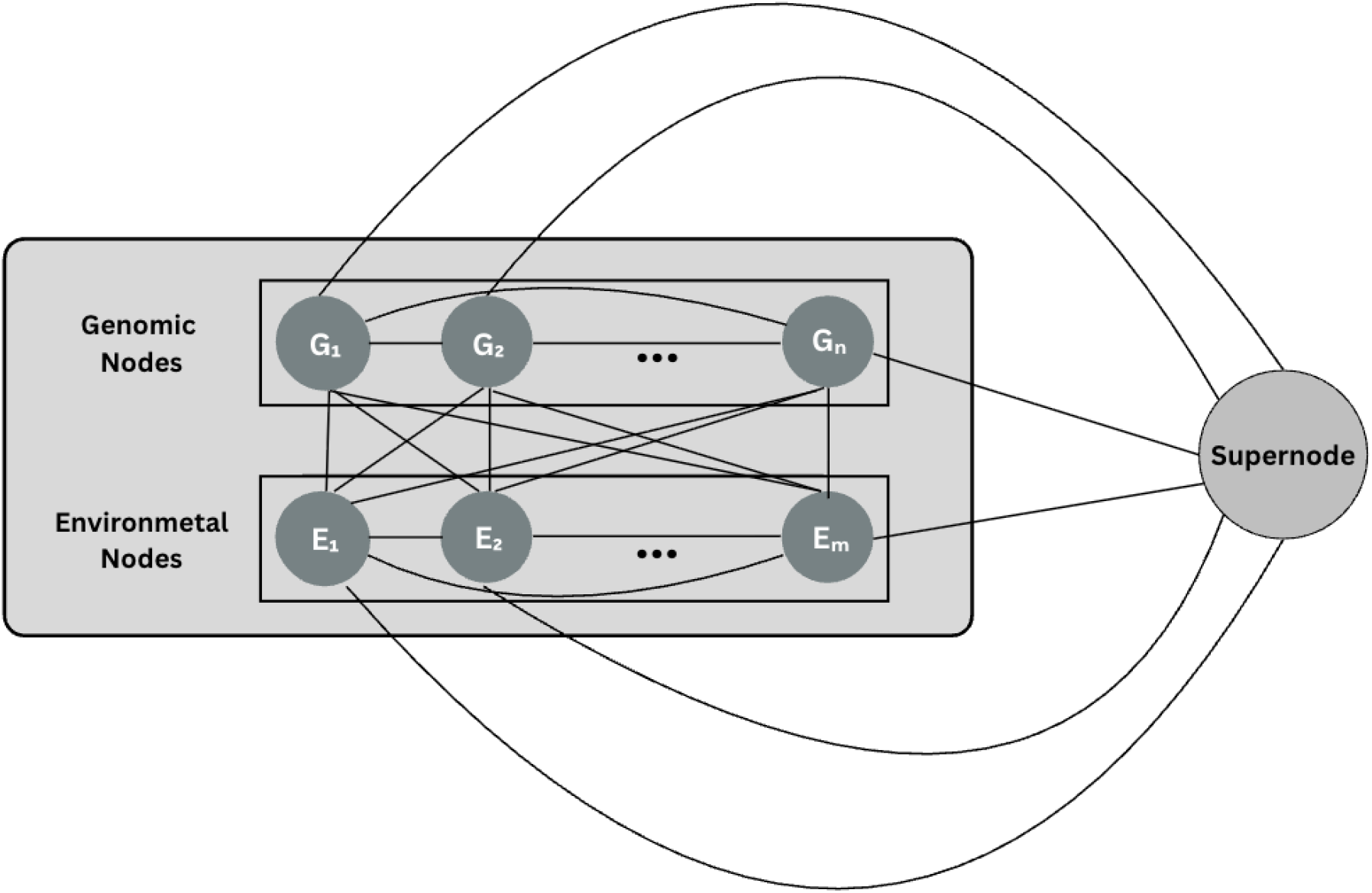
Detailed illustration of Architecture C showing the bipartite genotype–environment structure augmented with intra-genotype and intra-environment similarity edges with supernode.

Let 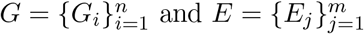 represent genotype and environment nodes, respectively. To create devoted directed top-*k* intra-set adjacencies *A*_*GG*_ and *A*_*EE*_ we use equation 14, and keep all bipartite edges *A*_*GE*_ and *A*_*EG*_.

After *K* iterations of message passing, we obtain updated embeddings for the *n* genotype nodes and *m* environment nodes, denoted by 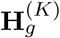 and 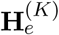. With the help of multi-head attention, the supernode query *s* conveys content-dependent weights to each node, allowing the model to highlight the most informative genotype and environment signals. Formally, the attention weights are defined as:

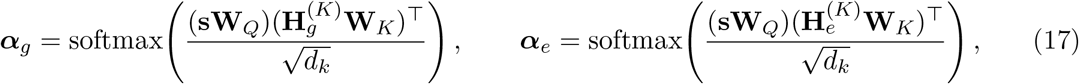

and the weighted sums yield

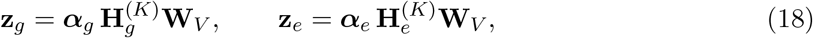

where **W**_*Q*_, **W**_*K*_, **W**_*V*_ are projection matrices and *d*_*k*_ is the key/query dimension. Finally, the two summaries are concatenated to form

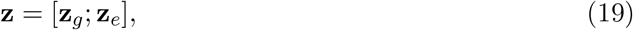

which serves as a compact representation of the genotype–environment graph and is passed to the MLP predictor.

The supernode readout is global and can attend to all nodes while remaining content-adaptive, as the weights depend on the learned attention mechanism. The supernode attention is executed after all message-passing layers have finished; that is, it operates on the final node embeddings that already incorporate genotype–environment context. This ensures that the global summary captures both local neighborhood effects and higher-order multi-hop relationships that were propagated through the GNN.

### 3.9 Hyper Parameter Tuning

We tuned all architectures under the same protocol described in Secs. 3.8.1–3.8.3: forward-time validation (2014–2021 train, 2022 test). The hyper parameters are described in Table 1.

**Table 1:** Final training configuration across architectures. A uses only bipartite edges (no intraset *k*); B adds intra-set edges; C keeps B’s message passing but replaces readout with a single global supernode attention pooling computed *after* message passing.

| Hyperparameter | Arch A | Arch B | Arch C |
| --- | --- | --- | --- |
| Batch size | 32 | 32 | 32 |
| Epochs | 100 | 100 | 100 |
| Intra-set $k$ ($A_{GG}, A_{EE}$) | — (not used) | 10 | 10 |
| Hidden dim $d$ | 128 | 128 | 128 |
| Attention heads | 8 | 8 | 8 |
| Learning rate | $3 \times 10^{-4}$ | $3 \times 10^{-4}$ | $3 \times 10^{-4}$ |
| Weight decay | $1 \times 10^{-4}$ | $1 \times 10^{-4}$ | $1 \times 10^{-4}$ |
| Workers | 4 | 4 | 4 |
| MLP (3-layer architecture) | 256–128–64–1 | 256–128–64–1 | 256–128–64–1 |
| Optimizer | AdamW | AdamW | AdamW |
| Loss | $\lambda \text{MSE} + (1-\lambda) \text{MAE}$ | $\lambda \text{MSE} + (1-\lambda) \text{MAE}$ | $\lambda \text{MSE} + (1-\lambda) \text{MAE}$ |

## 4 Results

### 4.1 Architectural Comparison

We evaluated the performance of three GNN designs that were trained under the same data splits and training conditions: A (fully bipartite), B (bipartite + intra-set edges), and C (the same message passing as in B but with a global supernode readout applied after the message passing). The performance was evaluated using RMSE, PCC, and *R*^2^ over the unseen gentypes and unseen environments as a test-set.

For the three architectures A, B and C under consideration, the increasing tendency can be seen in the performance with regard to accuracy and correlation (Table 2 and Figure 6). A, posts RMSE= 2.7749 and PCC= 0.4115. B, the error reduces to RMSE= 2.3683 and the PCC is boosted to 0.6622 which is 14.7% less error and 0.25 higher PCC than A. C, attains RMSE= 2.2120 and PCC= 0.6945 which is 20.3% less error than A and 6.6% less than B, with the highest correlation overall. The scatter plots for A, B and C show the strongest relationship between actual and predicted yield for C.

**Table 2:**
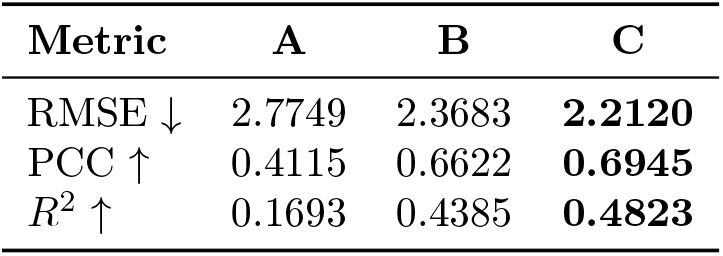
Test-set metrics across architectures.

**Figure 6:**
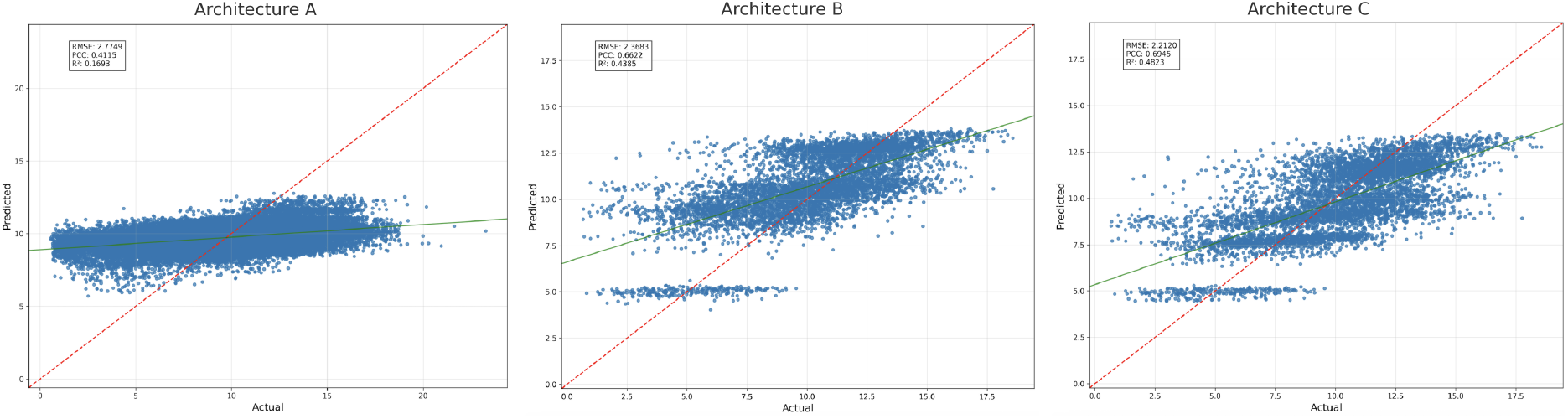
Predicted vs. actual yield on the test set for Architectures A, B, and C (left to right). The red dashed line is the identity; the green line is the fitted regression. Each point in the plot refers to an individual gentotype-site-year combination in the test set.

The steady progress of all three models in the training curves shown in Figure 7 is very evident. A shows a gradual improvement in both RMSE and PCC as the epochs progress. B obtains quicker results in PCC, however it has to sacrifice a little stability in RMSE during training. C has the clearest stage: an improvement phase during epochs ∼15–25 followed by a stable plateau at lower RMSE and higher PCC.

**Figure 7:**
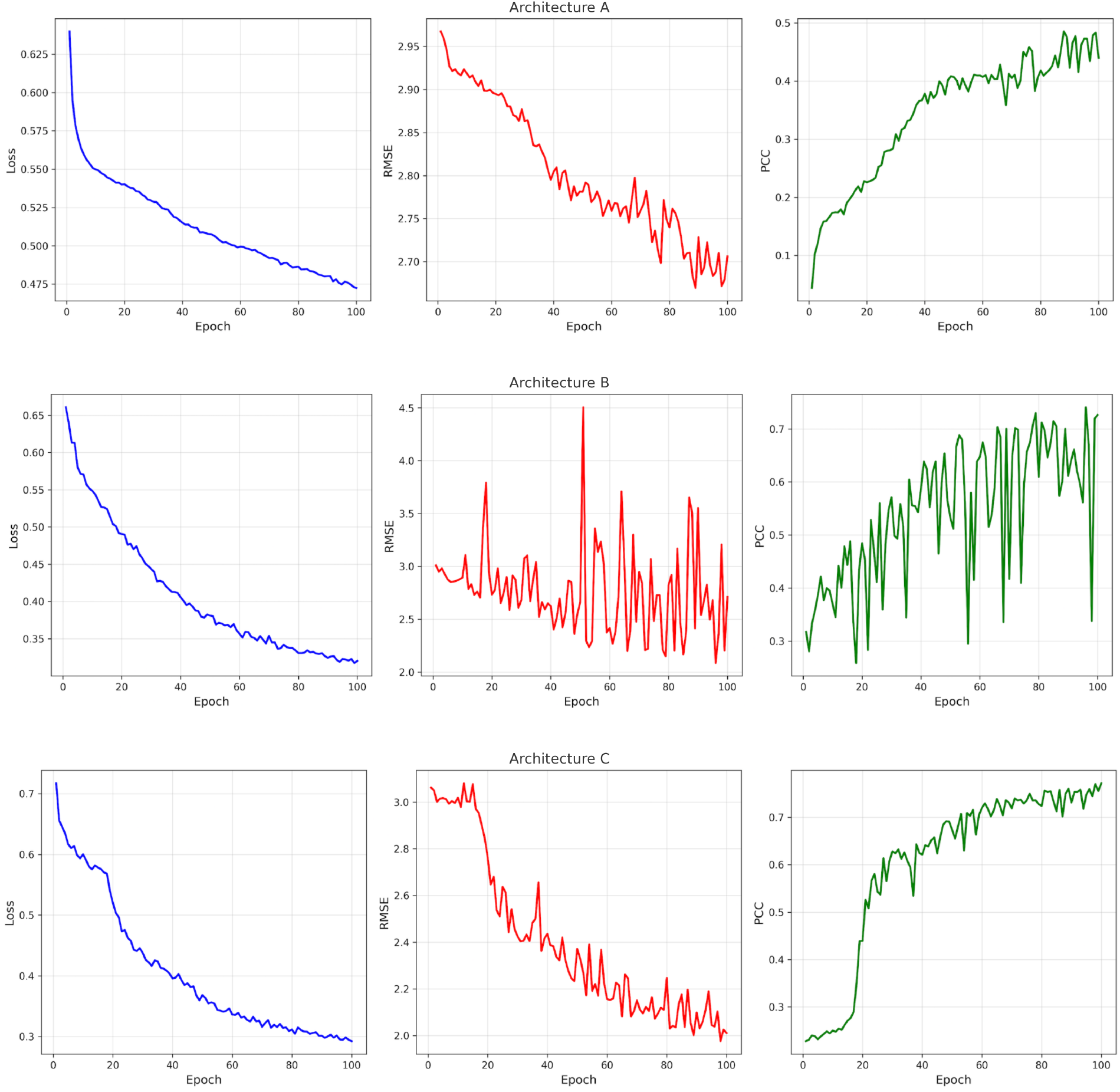
Training dynamics for A (top), B (middle), and C (bottom). Each row shows training loss (left), RMSE (middle), and PCC (right) across epochs.

### 4.2 Predictive Correlation vs. Genotype Count Across Site-Year

We used Architecture C, and performed the evaluation across all environments in the test set, which included genotype counts ranging from 335 to 530 (Figure 8). Although some environments show variability in PCC, the fitted regression line indicating no strong association between sample size in the environment and predictive accuracy. Thus, the model’s environment-level predictive ability appears independent of genotype count; increasing the number of genotypes in an environment (without changing covariates) does not automatically yield a higher PCC. Moreover, environments with larger numbers of genotypes are not influencing the system at the expense of those with fewer genotypes.

**Figure 8:**
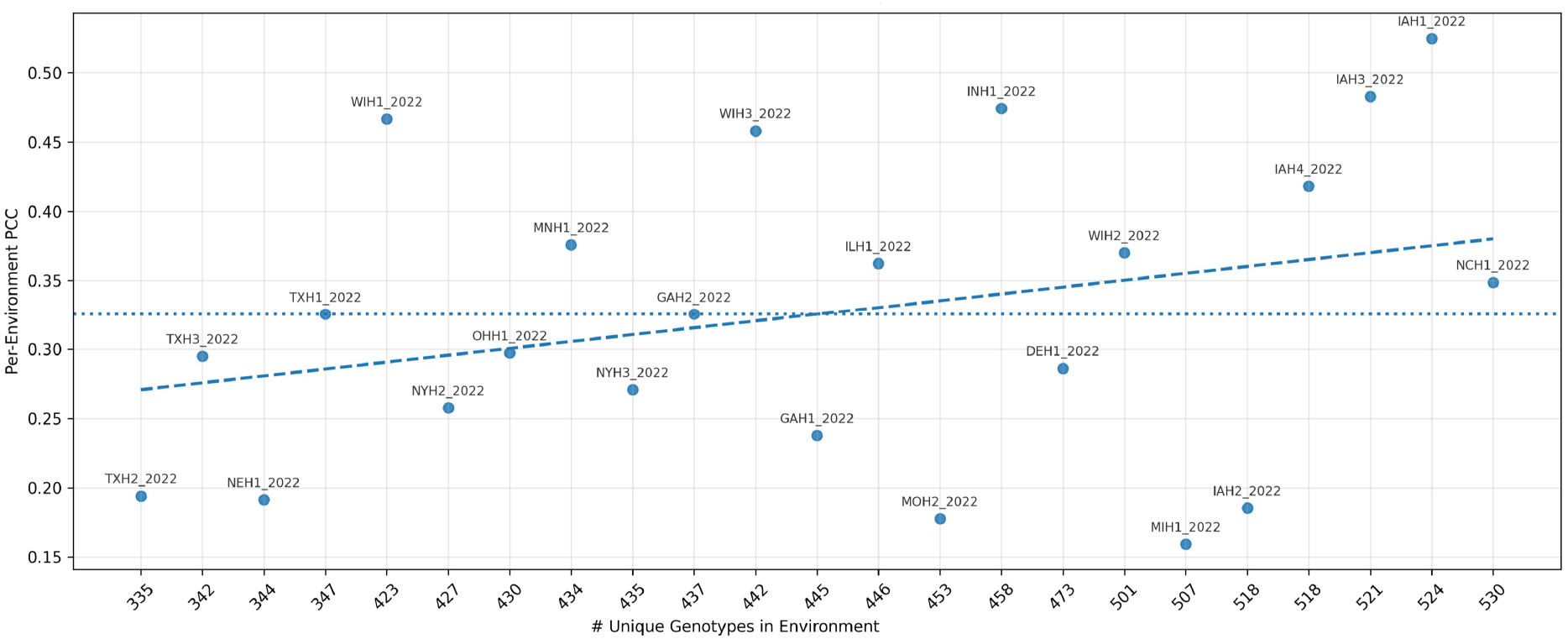
Per-environment predictive correlation (PCC) versus the number of unique genotypes evaluated.

### 4.3 Benchmarking against the G2F G*×*E Competition

We examine our method on the identical Genomes to Fields (G2F) dataset [9] and split utilized in the public G × E yield-prediction competition (2014–2021 training; 2022 held-out test). The competitive leader-board is based on the average RMSE values and Pearson correlation (PCC) of different environments. As a transparent external baseline, Table 3 shows our three models (A–C) with the top competition results.

**Table 3:** Comparison with the Global G*×*E Prediction Competition.

| Model | RMSE ↓ | PCC ↑ |
| --- | --- | --- |
| CLAC (winner) | 2.458 | 0.631 |
| DataJanitors (best PCC) | 2.752 | 0.644 |
| Architecture A | 2.7749 | 0.4115 |
| Architecture B | 2.3683 | 0.6622 |
| Architecture C (ours, best) | **2.2120** | **0.6945** |

The best model we have at present, which is Model C, has achieved RMSE = 2.2120 and PCC = 0.6945, thus, clearly surpassing the winner of the competition (CLAC) who obtained across-environment RMSE = 2.458; PCC = 0.631 and was the best reported across-environment PCC (DataJanitors; PCC = 0.644).

## 5 Discussion

Our findings indicate that graph neural networks are capable of capturing G × E interactions. By representing genotypes and environments as nodes within a graph, the GNN can model the complex dependencies between them. Consequently, the GNN was able to identify the G × E patterns, leading to improved yield predictions in our experiments.

The combination of local and global graph edges was one of the key factors that contributed to the success of the model in modeling G × E interactions. We incorporated intra-set edges among genomic and environmental nodes, which allows the model to exchange information between connected nodes and exploit similarities among related nodes within both domains.

We added a supernode that connects to every other node in the graph, helping the model learn global relationships. Through this design, the model captures higher-order similarities that exist among all genotypes or environments. Together, the local and global graph structures allow the model to better capture G*×*E interactions.

We used an LSTM to encode environmental time series data. Instead of a standard description of environmental features based on static summary statistics, we presented the sequential daily weather data (temperatures, precipitation, etc.) to an LSTM resulting in the condensed version of the growing season. The primary idea is that timing and duration of environmental events may influence crop yield in ways that cannot be explained by average conditions alone. This means that, an LSTM might learn that a late-season drought after an early wet period affects yield differently than an equally dry period occurring in mid-season.

Incorporating local subgraphs among genomic and environmental nodes increases the model’s complexity and allows better information exchange. These subgraphs help capture interactions within the genomic and environmental nodes, allowing them to share information within their groups and help the entire of the architecture to better model G*×*E interactions.

## 6 Conclusion

We proposed a GNN+LSTM joint model for G × E yield prediction. The GNN is responsible for capturing non-linear genotype-environment interactions through message passing, and the LSTM is responsible for transforming raw weather time series into compact environment embeddings. Two critical design approaches were local links (among genotypes and among environments) and a global connection (the supernode readout after message passing). The combination of these improved correlation and calibration.

We believe that future work in this direction can explore alternative data representations, such as constructing graphs in which each node corresponds to a genotype–environment pair, or clustering environments into broader regimes to evaluate genotype performance within and across these regimes.

## Declarations

### Funding

This work was supported by Natural Sciences and Engineering Research Council Discovery Grant to Mike Domaratzki.

### Conflict of interest

The authors declare no competing interests.

### Ethics approval

Not applicable.

### Consent for publication

Not applicable.

### Data availability

No data was created as part of this project.

### Code availability

Code will be made available upon publication.

### Author contribution

Amir Morshedian: Conceptualization, Methodology, Investigation, Writing Original Draft. Mike Domaratzki: Conceptualization, Supervision, Writing Review and Editing.

## Notes

### Competing Interest Statement

The authors have declared no competing interest.

